# Tunicate-specific protein Epi-1 is essential for conferring hydrophilicity to the larval tunic in the ascidian *Ciona*

**DOI:** 10.1101/2024.09.30.615758

**Authors:** Kazu Kuroiwa, Kaoru Mita-Yoshida, Mayuko Hamada, Akiko Hozumi, Atsuo S. Nishino, Yasunori Sasakura

## Abstract

Animals must avoid adhesion to objects in the environment to maintain their mobility and independence. The marine invertebrate chordate ascidians are characterized by an acellular matrix tunic enveloping their entire body for protection and swimming. The tunic of ascidian larvae consists of a surface cuticle layer and inner matrix layer. Hydrophilic substances coat the cuticle; this modification is thought to be for preventing adhesion. However, the molecule responsible for regulating this modification has not been clarified. We here found that the tunicate-specific protein Epi-1 is responsible for preventing adhesiveness of the tunic in the ascidian *Ciona intestinalis* Type A. *Ciona* mutants with homozygous knockouts of *Epi-1* exhibited adhesion to plastic plates and to other individuals. The cuticle of the *Epi-1* mutants was fragile, and it lost the glycosaminoglycans supplied by test cells, the accessory cells that normally attach to the tunic surface. Although it has an apparent signal peptide for membrane trafficking, we showed that the Epi-1 protein is localized to the cytosol of the epidermal cells. Our study demonstrated that the emergence of the tunicate-specific protein Epi-1 accelerated the immediate ancestor of tunicates to evolve a tunic by providing a way to avoid the adhesiveness of this structure.

**Highlights:**

- The ascidian *Ciona* tunic has a glycosaminoglycan (GAG)-coated hydrophilic cuticle.
- Epi-1 is a protein expressed in the epidermis of tunicates during embryogenesis.
- *Epi-1 Ciona* mutants have GAG-free, hydrophobic and therefore sticky cuticles.
- Epi-1 acquisition may have prevented adhesion of tunicates to environmental objects.

## Introduction

Tunicates are marine invertebrate chordates and the closest living relatives to vertebrates (Delsuc et al., 2006; Putnam et al., 2008). Their defining characteristic is the tunic, an acellular matrix that covers their entire body (Satoh, 1994). Among chordates, only tunicates possess the tunic. The shapes and functions of tunics vary among three tunicate subgroups: ascidians, salps, and larvaceans. The ascidians form the largest subgroup among tunicates, and their adults are sessile animals that metamorphose from swimming, tadpole-shaped larvae (Cloney, 1982; Karaiskou et al., 2015). The primary role of ascidian tunics is likely to be protection from predators. Moreover, at the larval stage, the tunic of ascidians forms the tail fin, which is suggested to efficiently capture the seawater to propel the body by tail beating (Cloney and Cavey, 1982). The larvaceans remain free-living animals throughout their life. The tunic of larvaceans forms the house, an extra-organismal apparatus that includes a net to assist in capturing foods in the seawater for efficient feeding (Thompson et al., 2001). The different uses of the tunic among the subgroups suggest that acquiring this structure was essential for the evolution and radiation of tunicates.

The larval tunic of the ascidians is divided into an inner matrix layer and a surface cuticle (Fig. 1). The matrix is characterized by fibrous cellulose secreted from the epidermis (Nakashima et al., 2004; Ranby, 1952; Sasakura et al., 2005). The cuticle is recognizable under an electron microscope as a series of thin electron-dense matrices (Hirose et al., 1997). Many species in the suborder Phlebobranchia of the order Enterogona, including the model tunicate *Ciona intestinalis* Type A (for which a renaming to *Ciona robusta* was recently suggested (Brunetti et al., 2015)), have smooth cuticles; however, the cuticles of some species in the suborder Aplousobranchia and many species in the order Pleurogona have multiple protrusions that are 20-130 nm high (Hirose et al., 1997). At the larval stage, test cells, maternally derived cells located at the perivitelline cavity between the chorion and the outer surface of the egg and embryo before hatching, attach to the surface of the tunic. The test cells have been suggested to play a role in the tunic formation (Cloney, 1990; Cloney and Hansson, 1996; Sato and Morisawa, 1999).

**Fig. 1.**
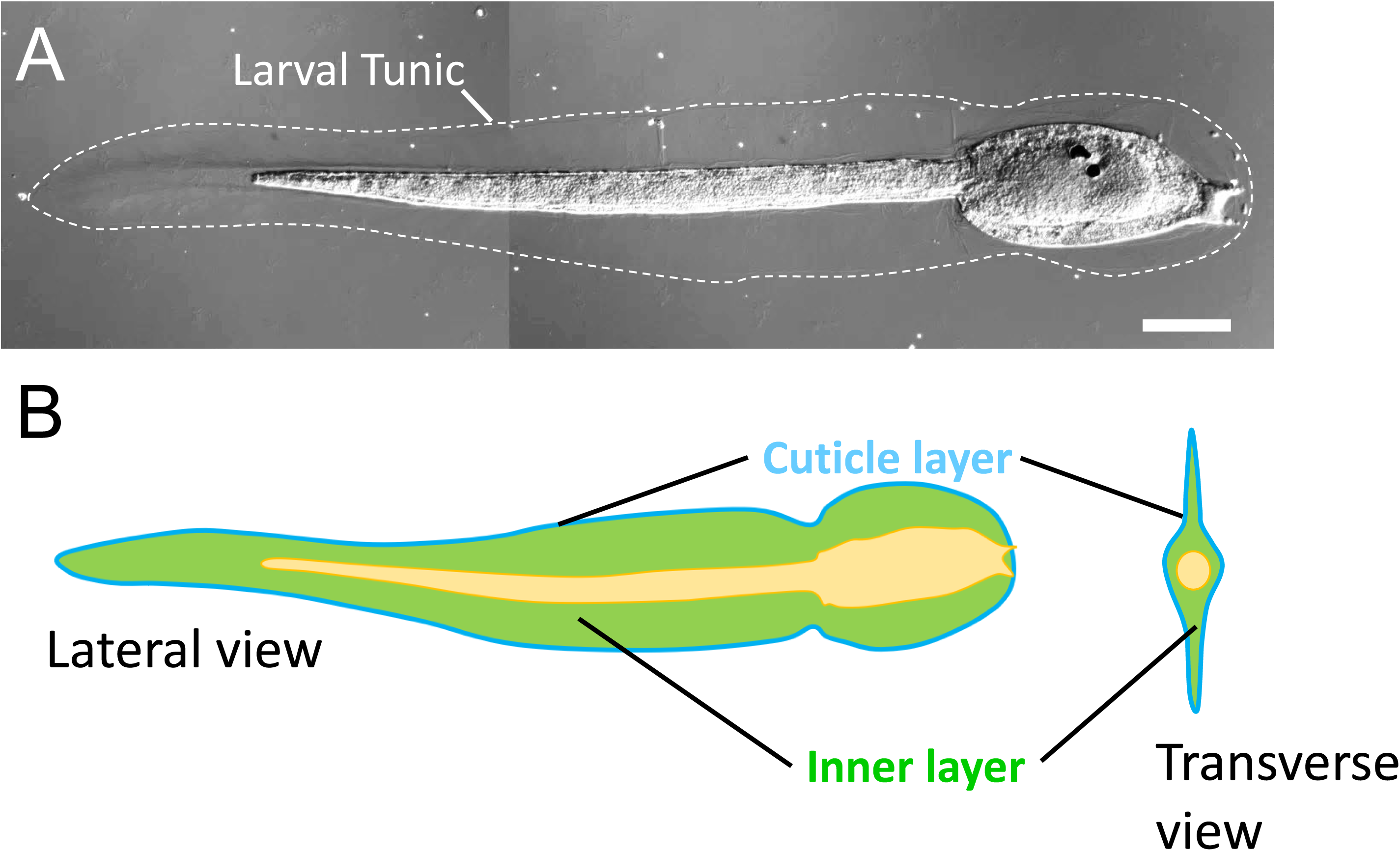
The organization of the larval tunic. (A) A tadpole larva of *Ciona intestinalis* Type A, lateral view. The anterior is toward the right, and the dorsal is toward the top. The larval tunic is outlined by a dotted line. (B) A schematic illustrating the layers of the larval tunic. Please note that *Ciona* larva possess an adult tunic near the larval body, which is omitted from the illustration for simplicity.

Cellulose production is a unique characteristic of tunicates. Because cellulose is a main component of the tunic, the acquisition of cellulose-producing ability was likely to have promoted the ancestor of tunicates to acquire a tunic. However, this event would not have been sufficient to obtain a functional tunic (Sasakura et al., 2005). Organisms must avoid adhesion to environmental objects such as other organisms, individuals of their own species, or inorganic substances. This prevention is necessary to maintain their identities, mobility, and survivability. In the case of ascidian larvae, these animals were suggested to avoid adhesion by masking the hydrophobic properties of the tunic (Cloney and Hansson, 1996). Test cells provide polysaccharides such as glycosaminoglycans (GAGs) to the tunic surface (Sato and Morisawa, 1999). Because GAGs are hydrophilic, this modification could mask the hydrophobicity of the tunic. Hydrophilicity can prevent object adhesions under aquatic conditions, suggesting that the acquisition of this mechanism enabled the ancestor of ascidians (or tunicates) to avoid adhesion to objects to maintain their freedom of movement. Therefore, we need to characterize the molecules responsible for controlling tunic hydrophilicity/hydrophobicity to deepen our understanding of the evolutionary history of tunic acquisition.

The acquisition of a novel trait is sometimes associated with acquiring novel proteins and genes. For example, the cellulose-producing ability of tunicates emerged by acquiring the gene encoding cellulose synthase from bacteria through horizontal gene transfer (Matthysse et al., 2004; Nakashima et al., 2004; Sagane et al., 2010; Sasakura et al., 2005). Because the loss of cellulose does not result in the abolishment of the tunic, additional genes are likely to be involved in tunic formation (Lanoizelet et al., 2024; Li et al., 2023; Sasakura et al., 2005).

Epi-1 is a protein first reported in *Ciona savignyi* as a marker of epidermis during embryogenesis (Chiba et al., 1998). Epi-1 has a signal peptide for membrane trafficking or secretion at its N-terminus and P-type trefoil domains that function in the protein–protein interactions. From the domain compositions, Epi-1 was predicted to be secreted from the epidermis to compose the mucus in the tunic. Although trefoil domain-containing proteins are encoded in vertebrate genomes, the clear orthologue of Epi-1 has not been found in animals other than tunicates, suggesting that Epi-1 is specific to tunicates and may have contributed to the acquisition of the tunicate-specific properties. However, the function of Epi-1 has not been fully addressed. In this study, we created and observed *Epi-1*-knockout mutants of *Ciona*. We found that mutant larvae with homozygous knockout of *Epi-1* have sticky tunics like those in larvae developed from dechorionated eggs. Our study demonstrates that the non-adhesive property of the tunic, a tunicate-specific structure, is achieved by a novel protein acquired specifically by this animal group.

## Results

### Epi-1 is necessary for maintaining the non-adhesive property of the larval tunic

To address the function of *Epi-1*, we knocked out this gene using the genome editing tool Transcription-Activator-Like Effector Nuclease (TALEN; Doyle et al., 2012; Sakuma et al., 2013; Treen et al., 2014). We constructed two pairs of TALENs (TAL1 and TAL2) that target the different nucleotide stretches at the 12th exon encoding the trefoil domain (Fig. 2). Both TALEN pairs efficiently introduced mutations in their target sites in the G0 generation (Fig. 2A and B). By screening the progeny derived from the sperm of the *Epi-1* TALEN-introduced G0 animals and wild-type eggs, we established *Epi-1^TAL1^* and *Epi-1^TAL2^* mutant lines. *Epi-1^TAL1^* and *Epi-1^TAL2^*, respectively, have 114 base pair (bp) and 5 bp deletions in exon 12, which are expected to cause truncation of the Epi-1 protein (Fig. 2C and D).

**Fig. 2.**
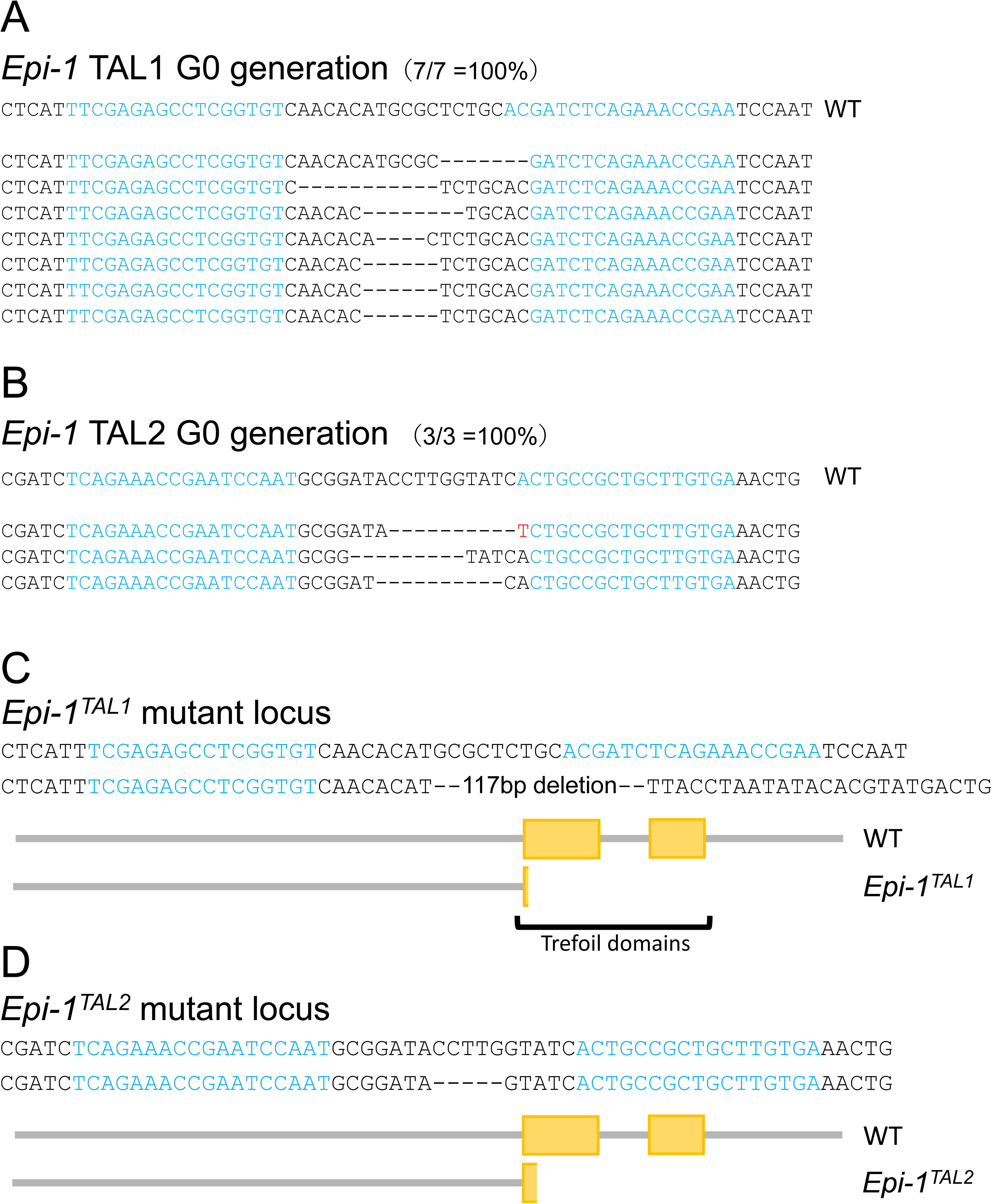
Epi-1 knockouts with TALENs. (A) Examples of the mutations induced by the pair of TAL1 constructs at the G0 (TALEN-introduced) generation. TALEN binding sites are indicated in blue. The numbers in the parentheses indicate the number of clones with mutations among the sequenced clones. Bars represent deleted nucleotides. WT represents the reference sequence at the target site. (B) Examples of the mutations induced by the pair of TAL2 constructs at the G0 generation. The red character indicates an inserted nucleotide. (C) The mutation harbored by the *Epi-1^TAL1^*mutant line, and the deduced protein structure produced by this mutated *Epi-1* locus. (D) The mutation harbored by the *Epi-1^TAL2^* mutant line.

The offspring larvae isolated by self-fertilization of a heterozygous *Epi-1^TAL1^* mutant included individuals without a notable fin along the dorsoventral axis (Fig. 3A and B). Moreover, they stuck onto the plastic dish and could not swim freely (Supplementary Movie S1 and S2). We concluded that the mutations in the *Epi-1* gene are causative of the adhesive phenotype. We observed 120 larvae, among which 32 individuals exhibited the adhesive phenotype. This score met Mendel’s law for a heterozygous locus (*p*<0.01, Chi-square test). Their genotyping exhibited that all sticky larvae were homozygous for the concerning *Epi-1^TAL1^* locus (Fig. 3D). In contrast, no homozygous *Epi-1* mutant was included in the wild types. 47 out of 200 offsprings isolated by self-fertilization of an *Epi-1^TAL2^* heterozygous animal exhibited the sticky phenotype (Fig. 3C; *p*<0.01, Chi-square test). The sticky larvae did not include a heterozygous animal (Fig. 3E), and their bands were homozygous for the *Epi-1* mutation. In the following experiments, we mainly used *Epi-1^TAL1^* for analyses. The *Epi-1* homozygous mutant larvae initiated metamorphosis and completed tail regression (Fig. 3F, 100%, n=45); however, they did not exhibit body growth and died within a few days.

**Fig. 3.**
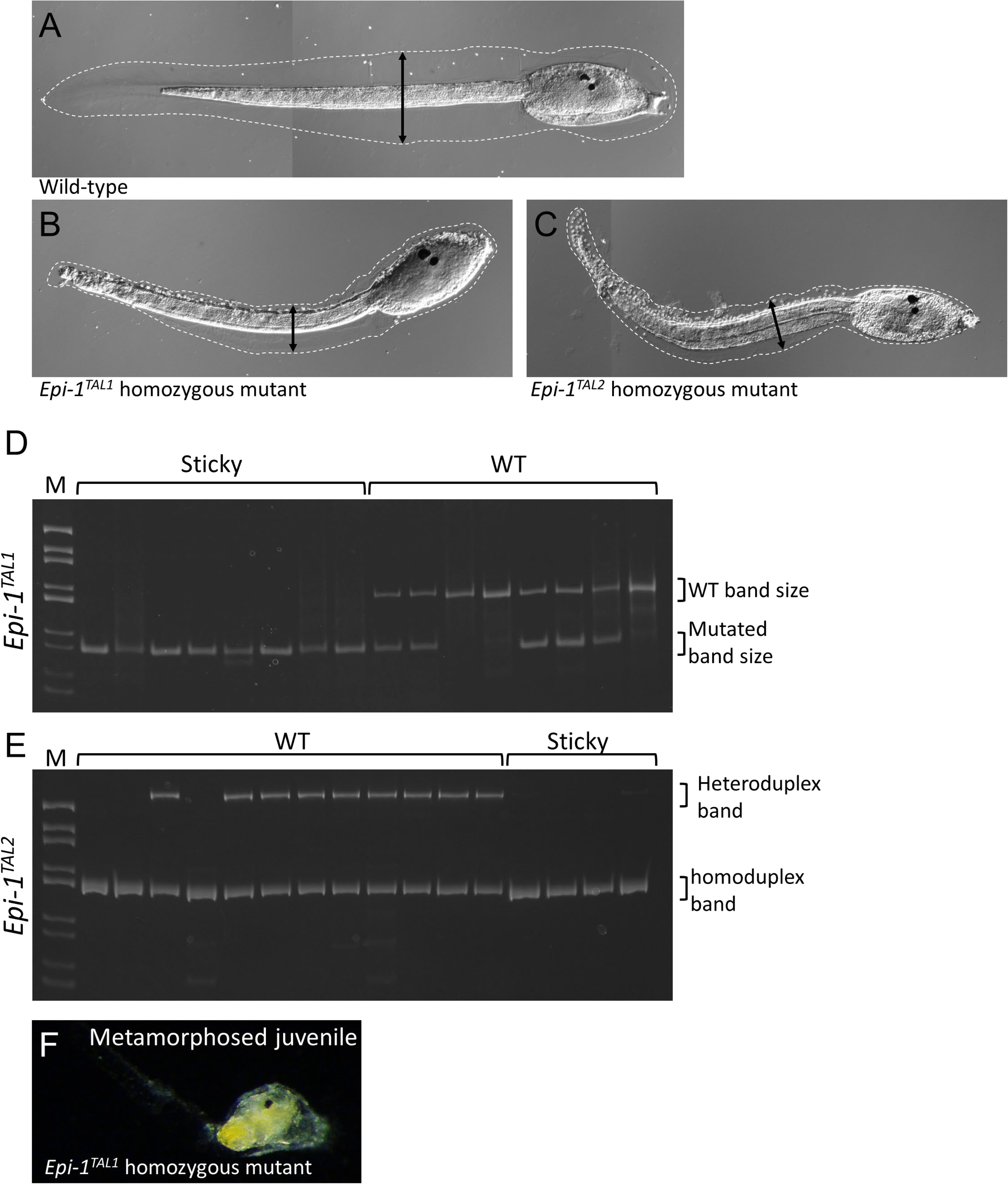
*Epi-1* mutants have sticky and malformed tunics. (A) A wild-type larva. Note that this panel is the same as Fig. 1A. The arrow represents the width of the tunic along the dorsoventral axis. (B) An *Epi-1^TAL1^*homozygous mutant larva with a narrow tunic. (C) An *Epi-1^TAL2^* homozygous mutant larva. (D) *Epi-1^TAL1^*homozygous mutant larvae had a sticky tunic. The DNA around the TALEN-targeted site was PCR-amplified and was subjected to electrophoresis. Because the *Epi-1^TAL1^* locus has a long deletion, the DNA from the locus yielded a shorter band that could be recognized by electrophoresis (Mutated band size). Each lane represents the result of the genotyping of a single larva, whose phenotypes are described above. WT, wild-type. M, marker lane (pBluescript digested by *Hae*III). (E) The genotyping of *Epi-1^TAL2^* mutants. This mutant locus has a 5 nucleotide deletion, which is too short to detect by electrophoresis. A heterozygous mutant yields a mixture of wild-type and mutant DNA. The heteroduplexes between the two DNAs have a structure different from homoduplexes, which resulted in the retarded mobility by electrophoresis (Heteroduplex band). Sticky larvae did not contain a heterozygous mutant, suggesting that they are homozygous for the *Epi-1^TAL2^* locus, which was confirmed by sequencing. (F) An *Epi-1^TAL1^*homozygous mutant after completion of tail regression.

### *Epi-1* mutants have incomplete cuticles and malformation of cellulose fiber

We observed the tunics of *Epi-1* homozygous mutant larvae using a scanning electron microscope (SEM) because the sticky phenotype of *Epi-1* mutants is likely to be associated with this outermost structure. The tunics of both the wild-type and *Epi-1* mutant larvae were covered by a cuticle that could be recognized as a smooth surfaced membrane (Fig. 4A-D). Test cells were adhered to the tunic surface in both wild-type and mutant larvae (Fig. 4B and D). However, the cuticle of *Epi-1* mutants was torn in several parts, and the fibers at the matrix layer were visible from the cracks (Fig. 4C, red rectangle, and 4E). These results suggest that *Epi-1* is not required for covering the tunic surface with the cuticle, but rather plays a role in providing stability to the cuticle.

**Fig. 4.**
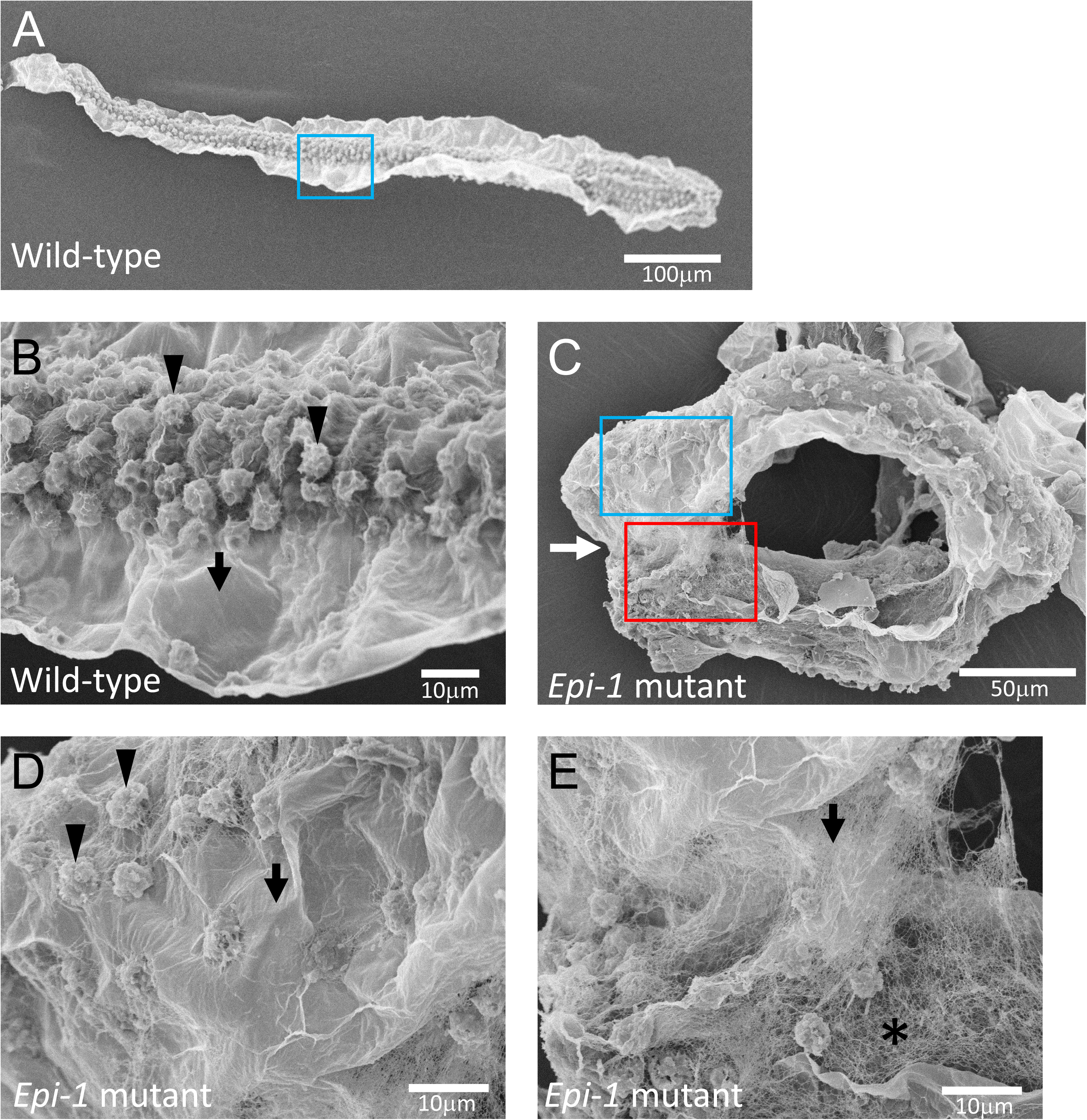
The larval tunic of the *Epi-1* mutants had a cuticle layer, as revealed by scanning electron microscopy (SEM). (A) An SEM image of a wild-type larva. (B) An enlarged image of the surface of the tunic in panel A (blue rectangle). Arrow, the smooth cuticle covering the matrix. Arrowheads indicate two examples of the tunic cells. (C) An SEM image of an *Epi-1* mutant larva. (D-E) Enlarged images of the tunic surface in panel C. Panels D and E show the area indicated by the blue and red rectangles, respectively. Arrowheads indicate two examples of the tunic cells. In E, an area exhibiting the fibrils in the mantle layer is shown by an asterisk.

The fibers in the mantle layer of *Epi-1* mutant larvae appeared irregularly twisted (Fig. 4E). These fibers were likely to be cellulose. To confirm this, we stained cellulose fibers in *Epi-1* mutants using GFP-CBM3, a fusion protein of green fluorescent protein and a cellulose-binding motif. The cellulose fibers in the tail of wild-type larvae were elongated toward the edge of the tunic to make up the fin shape (Fig. 5A-C). In contrast, the cellulose fibers in the *Epi-1* mutant were twisted toward random orientations (Fig. 5D and E), as the observation using SEM suggested. Therefore, we concluded that *Epi-1* is required to properly extend cellulose fibers in a unidirectional manner to form the fin.

**Fig. 5.**
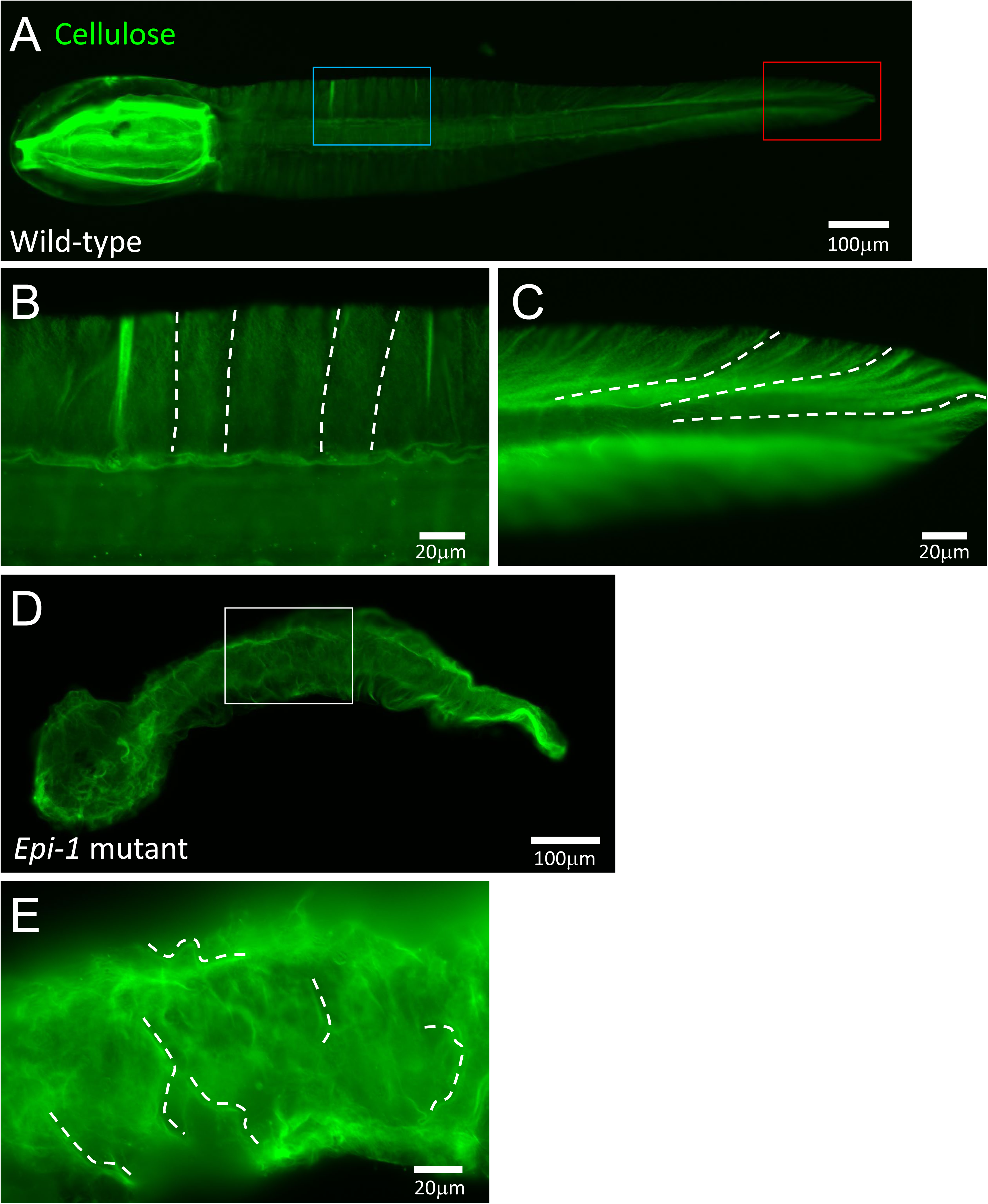
Cellulose fibrils in the *Epi-1* mutant tunic, shown in green pseudocolor. (A-C) A wild-type larva. (B) and (C) are the enlarged images of the middle and posterior parts, as shown in blue and red rectangles in panel A. The direction of cellulose fibrils is guided by the dotted lines. (D-E) An *Epi-1* mutant larva, and its enlarged image, showing the unorganized direction of cellulose fibrils.

### Epi-1 is a cytosolic protein

We next addressed the localization of the Epi-1 protein to elucidate its function in tunic formation. We made a polyclonal antibody to this protein that recognizes a sequence near its C-terminal end. This antibody bound specifically to Epi-1, based on the single band at around 80 kilodaltons (kDa) in the western blotting (Fig. 6A). The estimated size of the Epi-1 protein according to its primary structure is 80.8 kDa. Moreover, the western blotting of larvae that overexpressed the Epi-1::GFP fusion protein exhibited an additional higher molecular weight band (Fig. 6A) that was absent in the larvae overexpressing GFP in the epidermis (Fig. 6A). These data further confirmed that the antibody recognized the Epi-1 protein.

**Fig. 6.**
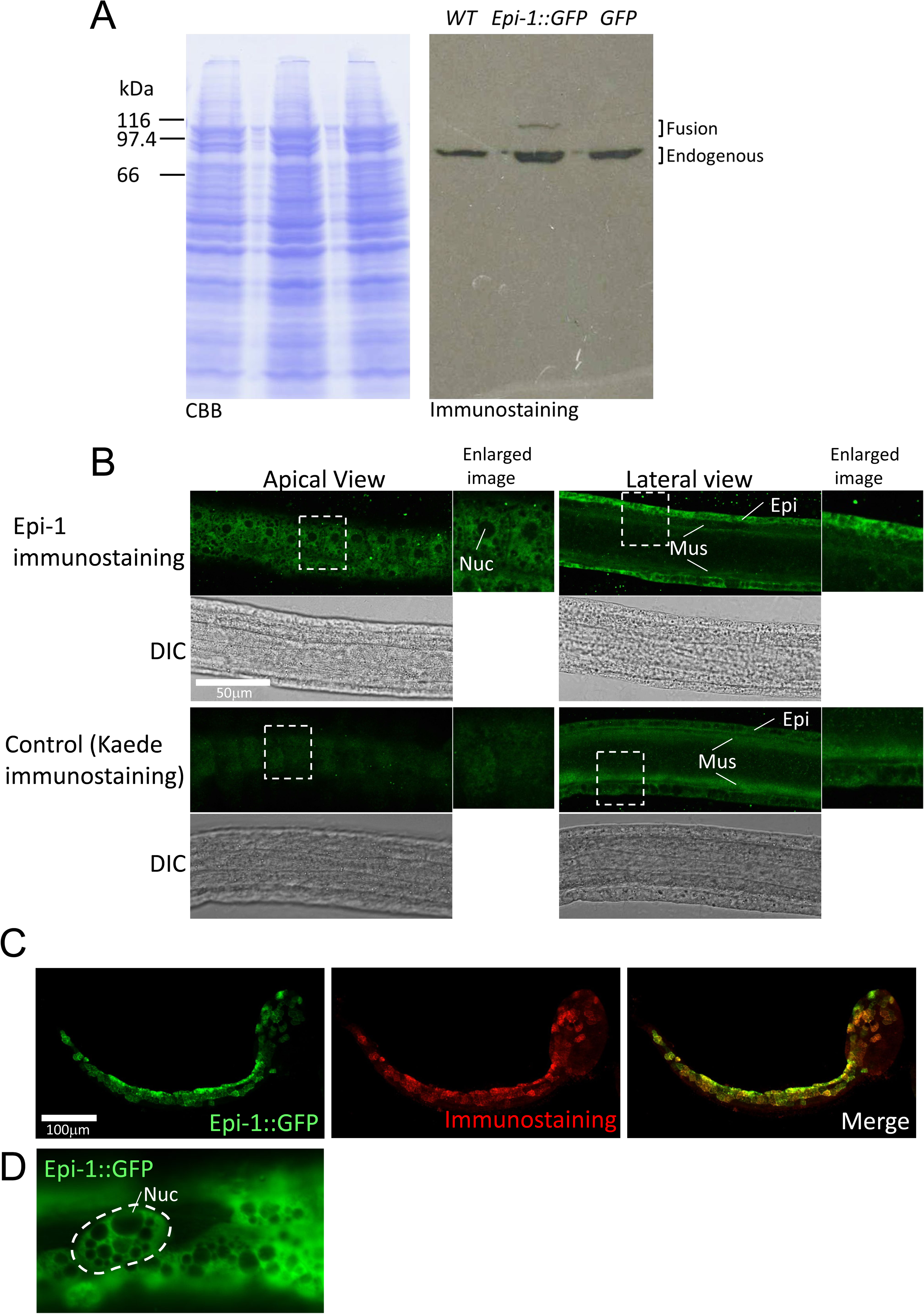
Epi-1 is a cytosolic protein. (A) A Western blot image of the Epi-1 protein. The left panel shows the Coomassie brilliant blue staining of the blotted proteins, and the right panel shows the immunostaining of Epi-1. WT, sample from wild-type larvae. Epi-1::GFP, sample from larvae overexpressing the Epi-1::GFP fusion protein. GFP, a sample from larvae overexpressing GFP. In the WT and GFP lanes, single bands at around 80 kDa can be seen, while the Epi-1::GFP lane exhibits a higher molecular-weight band representing the fusion protein. (B) Immunostaining of the Epi-1 protein in wild-type larvae. DIC, differential contrast images. The dotted lines indicate the areas shown in the enlarged image panels. Epi, epidermis; Mus, muscle; Nuc, nucleus. Epidermal cells of *Ciona* have multiple membranous structures, the interiors of which are immunonegative and therefore highlighted by black dots in the images. (C) Immunostaining of Epi-1 in the larvae overexpressing the Epi-1::GFP fusion protein in the epidermis. Green and red pseudocolors, respectively, represent the GFP fluorescence and immunostaining signals. (D) Epi-1::GFP fusion is localized in the cytosol. An epidermal cell is highlighted by a dotted line as an example.

Immunostaining of larvae using this antibody gave an immunopositive signal that was uniformly distributed throughout the entire cytosol of epidermal cells in both the apical and lateral views (Fig. 6B). We did not find an Epi-1-positive signal on the tunic compared to negative controls. These results suggest that Epi-1 is a cytosolic protein. To show the specificity of the immunostaining pattern of Epi-1, we immunostained the larvae overexpressing the Epi-1::GFP fusion protein in a mosaic fashion (Fig. 6C). Their immunostaining signal overlapped with GFP fluorescence, suggesting that the antibody specifically detects Epi-1. The localization of the Epi-1::GFP fusion protein, as detected by GFP fluorescence, was also cytosolic, and we did not detect any recognizable signal on the tunic (Fig. 6C and D). These data confirmed that Epi-1 is a cytosolic protein.

The cytosolic localization of the Epi-1 protein was curious because this protein has a typical signal peptide for a secreted or membrane protein at its N-terminus. Because Epi-1 does not have a transmembrane domain, this protein was suggested to be a secreted protein (Chiba et al., 1998). The expression of *Epi-1* is started at the initial tailbud stage (Chiba et al., 1998) that is prior to hatching. If Epi-1 is secreted before hatching, the Epi-1 protein should be accumulated in the perivitelline cavity. To examine this possibility, we compared the quantity of the Epi-1 protein between embryos with and without chorions by western blotting. However, their signals did not show a significant difference (Fig. 7). Therefore, Epi-1 was unlikely to be secreted during embryogenesis or at the larval stage.

**Fig. 7.**
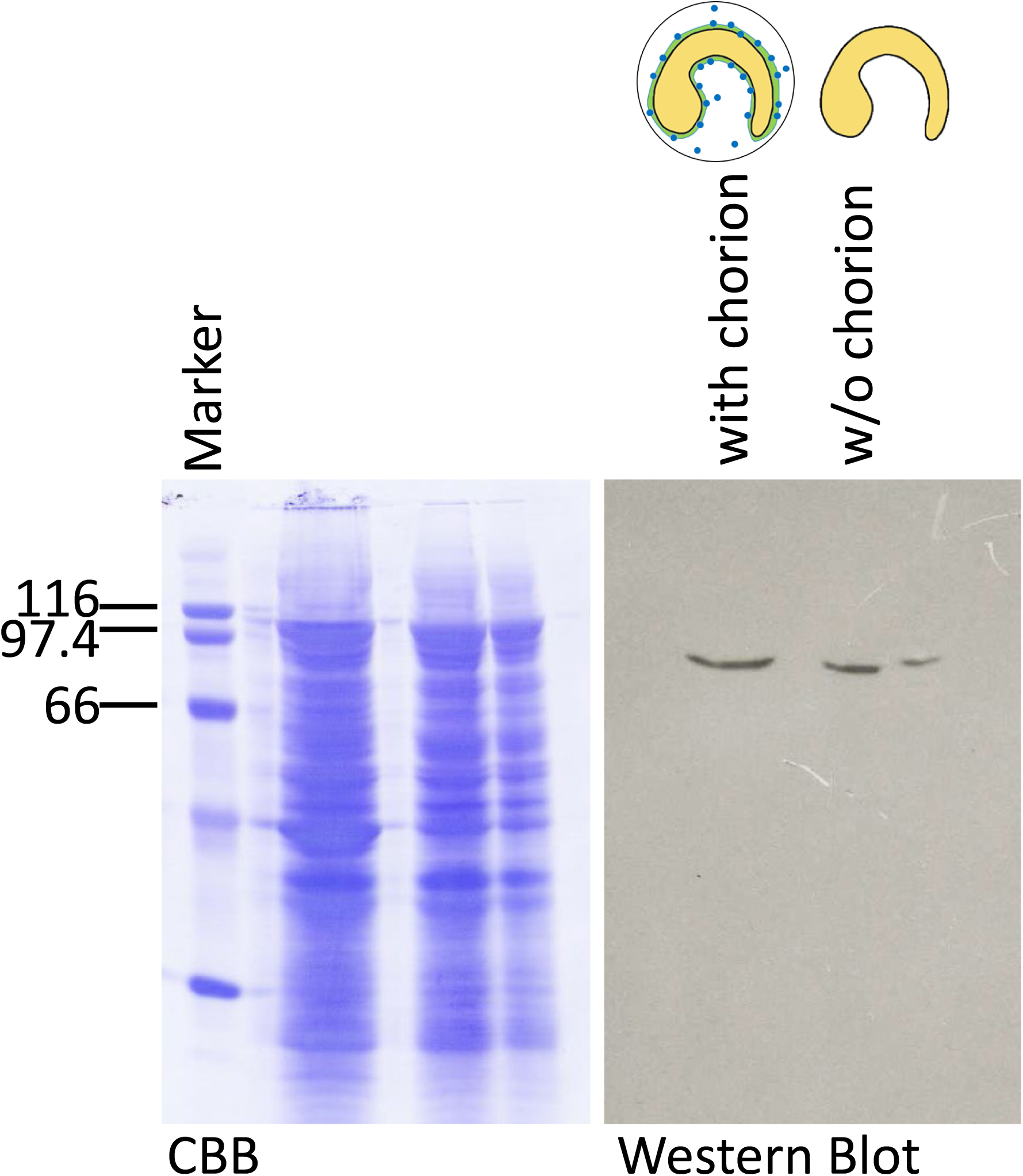
Epi-1 is not secreted into the perivitelline cavity. The signal intensities of the Epi-1 bands were compared between the samples with and those without chorions by immunoblotting. The proteins of the two lanes were derived from the same number of larvae. The two types of samples exhibited identical band intensities.

### *Epi-1* is required for covering the tunic surface with glycosaminoglycans secreted from test cells

The previous study showed that the surface of the larval tunic was glycosaminoglycan (GAG)-positive at successive time points after hatching, as detected by Alcian blue staining (Sato and Morisawa, 1999). The test cells were suggested to be the source of GAGs on the tunic surface because they became Alcian blue-negative when the tunic acquired GAGs. We stained *Epi-1* mutant larvae with Alcian blue to visualize their tunic surface properties. As a result, their larval tunics exhibited weaker staining of this dye compared to those of wild-type larvae (Figure 8A, A’, B, and B’). By contrast, the test cells of the *Epi-1* homozygous mutants exhibited stronger Alcian blue signals compared to those in wild-type larvae (Figure 8A’’ and B’’). Therefore, *Epi-1* is required for the test cells to secrete GAGs to cover the tunic surface.

**Fig. 8.**
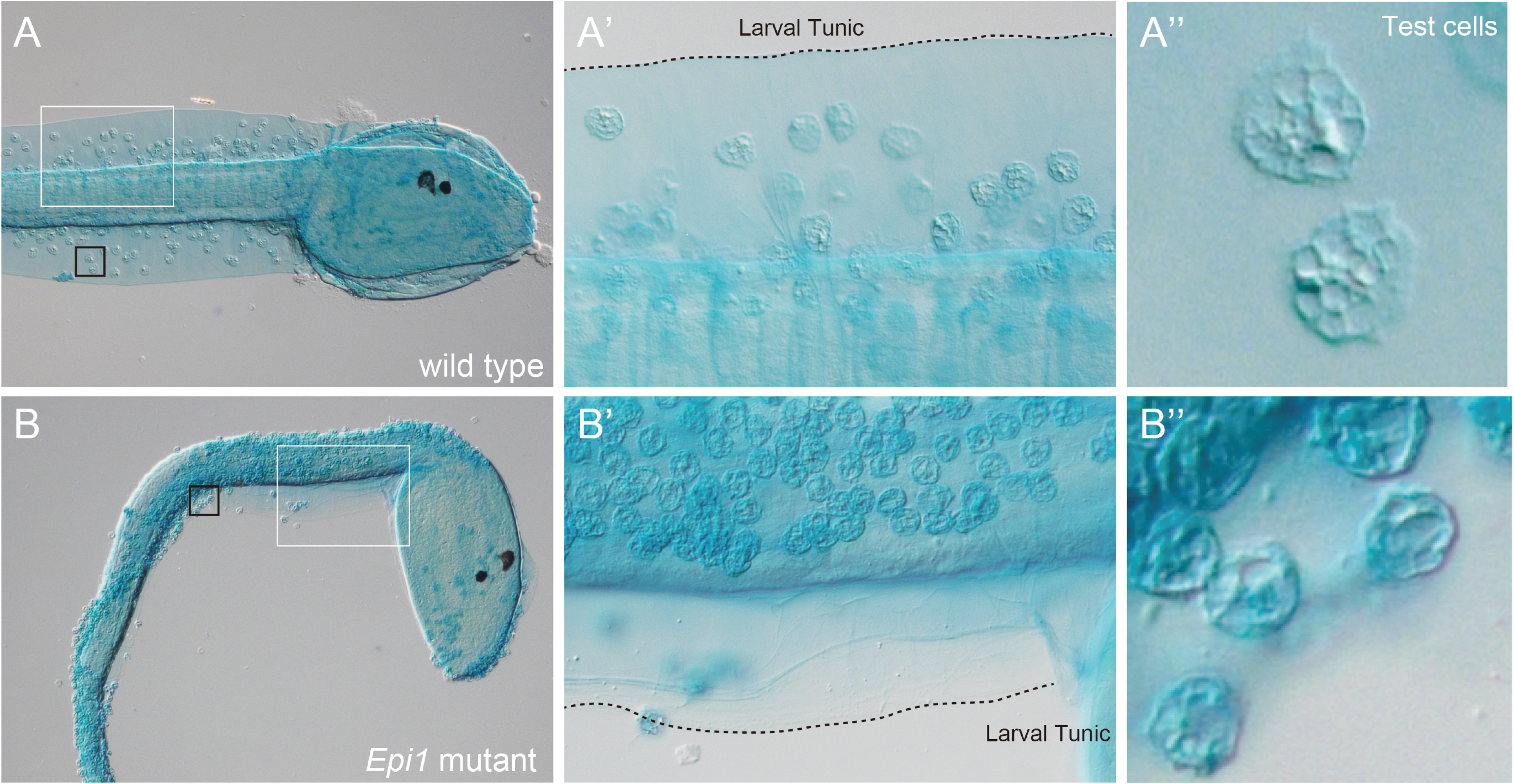
Glycosaminoglycans (GAGs) are absent on the surface of *Epi-1* mutant tunics. Blue represents GAGs stained by Alcian blue. (A-A’’) A wild-type larva. (A’) and (A’’) are the enlarged images of the regions shown with white and black lines in panel A, respectively. Dotted lines outline the larval tunic. (B-B’’) An *Epi-1* mutant larva.

We further examined the abnormality of the tunic surface of *Epi-1* mutants. Before metamorphosis, *Ciona* larvae secrete a mucus substance at the adhesive papillae. This substance, which could be detected using peanut agglutinin (PNA), was suggested to be an adhesive material that adheres larvae to a substrate (Zeng et al., 2019). We hypothesized that the sticky phenotype in *Epi-1* mutants is the ectopic secretion of this substance. However, PNA staining demonstrated that the adhesive was specifically secreted from the papillae of the homozygous mutants in *Epi-1* as was observed in wild types (Fig. 9A and B).

**Fig. 9.**
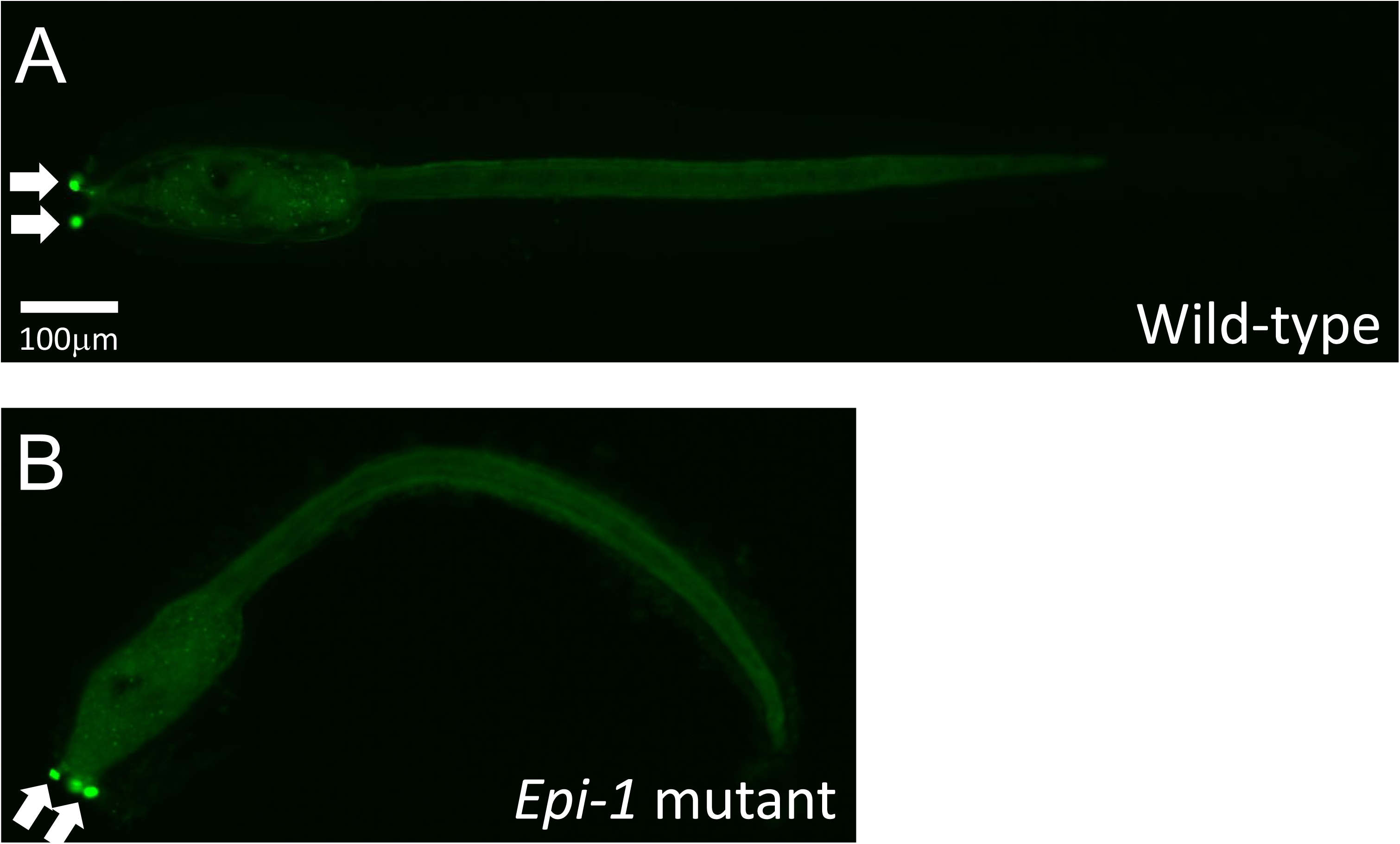
An adhesive substance is specifically secreted from the papillae of the *Epi-1* mutant, as revealed by PNA staining (green pseudocolor). (A) A wild-type larva. Arrows represent PNA-positive signals in the papillae. (B) An *Epi-1* mutant larva.

The cuticles of *Epi-1* mutant larvae had cracks through which cellulose fibers could contact external materials. To examine whether naked cellulose is the cause of their stickiness, we used the mutant of the gene encoding cellulose synthase (*CesA*; Sasakura et al., 2005) and a dechorionation protocol for the following reasons. If we had used double mutants of *CesA* and *Epi-1,* only one out of sixteen offspring larvae would be expected to be homozygous from a heterozygous mutant parent, which is too low a ratio to clearly verify the phenotype. The larvae developed from dechorionated eggs have undeveloped fins and sticky tunic surfaces (Cloney and Hansson, 1996), and thus it is highly likely that both *Epi-1* mutation and dechorionation influence the same processes forming the tunic. In addition, dechorionation can be applied to all larvae, which makes it easier to observe.

Homozygous mutants in *CesA* possess a malformed tunic that lacks cellulose fibers (Sasakura et al., 2005). Our observation using SEM confirmed that cellulose fibers are absent from the tunic of *CesA* mutant larvae, and their tunic is covered by the cuticle layer (Fig. 10A). The offspring from two *CesA* heterozygous mutants were dechorionated at the 1-cell stage. They were placed onto the culture dishes at the larval stage to see whether they adhered to the bottom. The results showed that all larvae adhered to the dish (Fig. 10B). We could not distinguish *CesA* homozygous larvae from the wild types after dechorionation because the abnormal tunic shape, the major phenotype of the *CesA* mutants one day after fertilization, is masked by the tunic malformation in the larvae developed from dechorionated eggs. However, the number of observed larvae statistically assures that the adhered larvae included *CesA* homozygous mutants, suggesting that naked cellulose is not causative of the sticky phenomenon in the *Epi-1* mutants and dechorionated larvae.

**Fig. 10.**
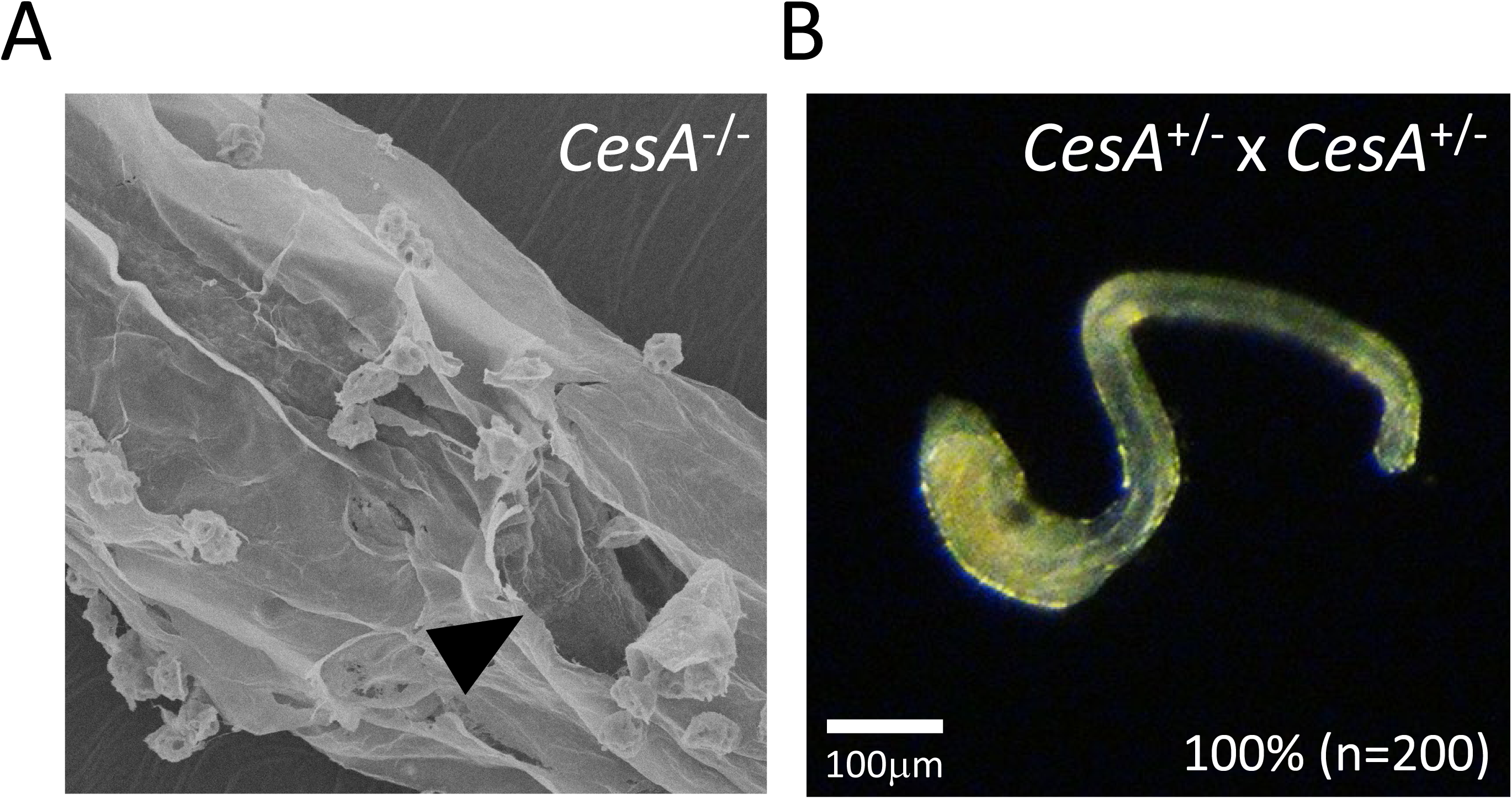
Naked cellulose fibrils were not the cause of the sticky tunic of the larvae under dechorionation conditions. (A) An SEM image of a *CesA* homozygous mutant larva. Fibers are absent in the mantle layer, as shown by the hole at the cuticle (arrowhead). (B) A sibling larva derived from two *CesA* heterozygous mutants. The siblings were developed under dechorionation conditions. This larva adhered to the plastic dish, and the body was twisted. The number at the bottom represents the number of larvae adhered to the dish.

## DISCUSSION

Epi-1 is a tunicate-specific protein found over 25 years ago (Chiba et al., 1998). We showed that this protein is required to prevent the larval tunic from adhesion to environmental objects. The tunic is a tunicate-specific structure that covers the entire body. The main component of the tunic is cellulose: the ancestor of tunicates acquired cellulose synthase from bacteria, which allowed them to form tunics (Matthysse et al., 2004; Nakashima et al., 2004). However, acquiring cellulose-producing ability was insufficient to make a functional tunic (Sasakura et al., 2005). Prevention of the adhesive property of the tunics is essential to ensure free-swimming activity during the larval stage and maintenance of solitary status at the adult stage. This study demonstrated that the emergence of *Epi-1* may have enabled the ancestor of tunicates to prevent this property of the tunic, having facilitated the use of the tunic as a novel structure in the tunicate lineage.

How does Epi-1 prevent adhesion? *Epi-1* mutant larvae and larvae from dechorionated eggs share the characteristic of sticky, glycosaminoglycan (GAG)-negative tunics. A previous study suggested that substances from test cells provide hydrophilicity to the tunic surface (Cloney and Hansson, 1996), and hydrophilic sugar chains are a strong candidate for such a substance (Sato and Morisawa, 1999). The loss of GAGs by the disruption of *Epi-1* and the loss of test cells by dechorionation is likely to make the tunic more hydrophobic, causing them to adhere to the plastics and other individuals. Indeed, coating plastic surfaces with hydrophilic proteins (such as albumin and gelatin) prevents larvae under dechorionated conditions from adhering to them (Yasuo and McDougall, 2018). Because the tunics of *Epi-1* mutants exhibit a cuticle layer with a smooth surface, which is visibly identical to the cuticles of wild-type larvae, the main component forming the cuticle surface is likely to have the hydrophobic property, which needs to be modified by the hydrophilic substances with the aid of Epi-1. Perhaps the smooth layer functions as a barrier to prevent the penetration of pathogens by tightly packing the components, and hydrophobicity is required for the tight packing.

Although Epi-1 has the signal peptide for secretion, our data suggests that Epi-1 is a cytosolic protein. As to the mechanism underlying this phenomenon, it may be that Epi-1 alters the nature of the proteins that build up the cuticle through protein– protein interactions using the trefoil domain in the epidermal cells (Fig. 11). Such modifications may not be essential for formation of the cuticle layer, but they are likely to function in the recognition between the tunic and test cells for promoting GAG secretion, and in providing rigidity to the cuticle. Because Epi-1 does not have a domain that directly modifies proteins, another protein may interact with Epi-1 and cuticle proteins. Likewise, the regulation of cellulose fibril formation by Epi-1 is likely to be indirect because the rosette terminal complex produces cellulose, and this production is the machinery of cellulose polymerization at the plasma membrane (Doblin et al., 2002; Kimura and Itoh, 2004). Because Epi-1 regulates the orientation of cellulose fibrils, Epi-1 may be a component of the rosette.

**Fig. 11.**
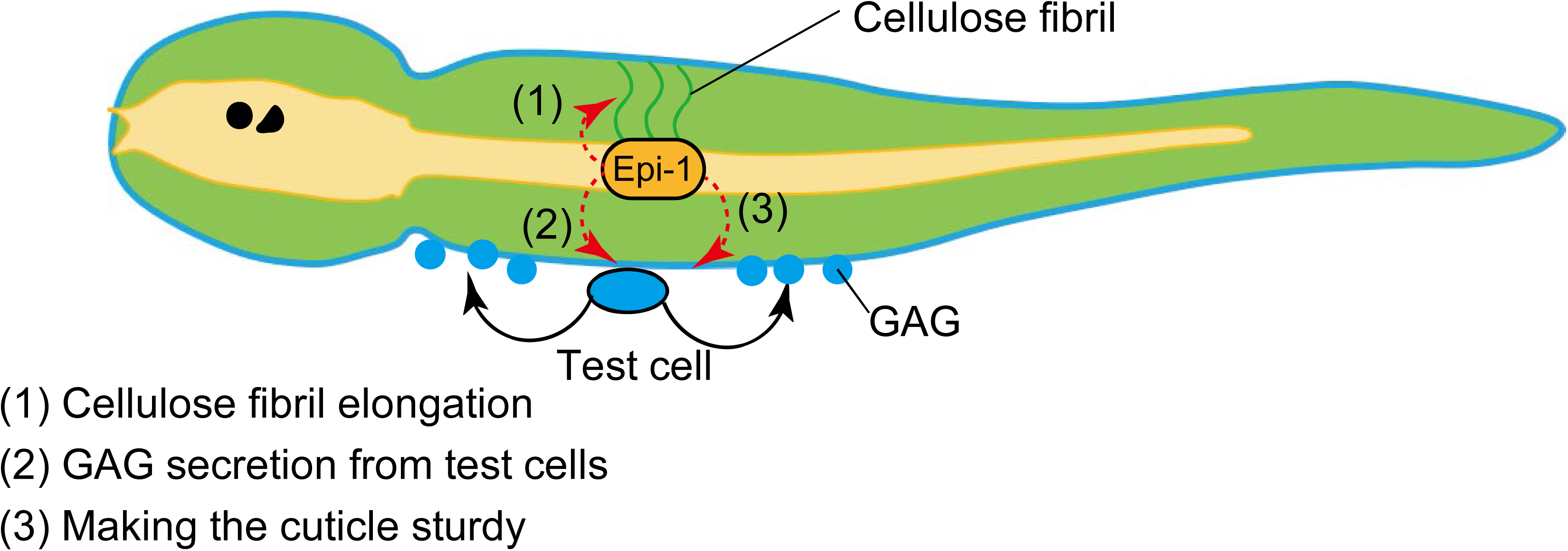
Schematic of the role of the Epi-1 protein in tunic formation. The oval represents an epidermal cell. Epi-1 in the epidermis mediates the cellulose fiber elongation, GAG secretion from the test cells, and the strength of the cuticle.

Acquiring novel traits is essential for evolution. The acquisition of new genes is a driving force for promoting this evolution. In tunicates, the tunic is the representative structure that supports fast swimming, efficient food uptake, and a sessile lifestyle. Acquisition of the cellulose-producing ability through horizontal gene transfer must have significantly promoted acquisition of the tunic (Matthysse et al., 2004; Nakashima et al., 2004; Sagane et al., 2010). In addition to the cellulose synthase gene, tunicates have multiple group-specific genes (Dehal et al., 2002). Some of them may function in establishing the tunic, as this study suggests. Although functional analyses of these tunicate-specific genes have been less actively pursued compared to functional analyses of the genes conserved among chordates, they must have been the lead players in promotion of the evolution of tunicates.

## Acknowledgments

We are grateful to the members of the Shimoda Marine Research Center at the University of Tsukuba for their maintenance of the animals and to Drs. Shigeki Fujiwara, Manabu Yoshida, and Yutaka Satou, and all members of the Department of Zoology, Kyoto University, the Misaki Marine Biological Station, the University of Tokyo, the Maizuru Fishery Research Station of Kyoto University and the National BioResource Project (NBRP) for the cultivation and provision of *Ciona* adults and experimental materials. We would also like to thank Drs. Ryuji Yanase and Kazuo Inaba for kindly instructing us on the microscopy using SEM. We are grateful to Dr. Nicholas Treen, Prof. Tetsushi Sakuma and Prof. Takashi Yamamoto for their support with the TALEN constructions. This study was supported by grants from the Japan Society for the Promotion of Science to YS (16H04815, 19H03262). YS was further supported by a Takeda Bioscience Research Grant.

## Data Availability

*Epi-1* mutants and plasmid constructs are available from the National BioResource Project, Japan. Requests for generated datasets should be directed to the corresponding authors.

## Author Contributions

KK, ASN, and YS conceived the project. KK performed most experiments with the assistance of AH, MH, and YS. AH carried out the immunostainings. KM carried out genetic analyses under consultation with MH. ASN supervised the experiments using larvae under dechorionated conditions and supported the design of this study. KK, ASN, and YS wrote the manuscript.

## Materials and Methods

### Animals

The National BioResource Project, Japan, cultivated wild-type *Ciona intestinalis* Type A/*Ciona robusta* individuals derived from Onagawa Bay (Miyagi, Japan) and Onahama Bay (Fukushima, Japan) as closed colonies. They were kept under constant light to avoid untimely spawning. Eggs and sperm were surgically collected from gonadal ducts. The *CesA* mutant was described previously (Sasakura et al., 2005). Mutants were maintained according to the in-land culture system (Joly et al., 2007). The recently modified conditions in the culturing were as follows: We exchanged seawater of 5L to 30L tanks once per week, and the condensed pastes of the algae *Pavlova sp.* and *Isochrysis sp.* (Reed Mariculture) were fed together with *Chaetoceros calcitrans*. Photographs were taken with AxioImager Z1 and AxioObserver Z1 (Carl Zeiss). Movies were taken with Handycam HDR-CX590 (Sony).

### Constructs

The TALEN pairs that target the region encoding the trefoil domain of *Epi-1* were constructed on the TALEN backbone vector having the *EF1α cis-*element and mCherry by the Golden Gate method based on previous reports (Cermak et al., 2011; Sakuma et al., 2013; Treen et al., 2014). The target sites of TALENs were determined using TAL Effector Nucleotide Target 2.0 (Doyle et al., 2012). The *EF1α cis-*element was switched to that of *CiNut* (Kitaura et al., 2007) with an In-Fusion Cloning Kit (Clontech) according to the previous reports (Sasakura et al., 2017; Yoshida et al., 2017; Yoshida and Treen, 2018). pSPCiEpi1>GFP (Sasakura et al., 2010) was subjected to inverse PCR with the primers 5’-GTGAGCAAGGGCGAGGAGCTG-3’ and 5’-CATTTTTTTGTATGCTACCAG-3’. The open reading frame of *Epi-1* was PCR amplified with the primers 5’-GCATACAAAAAAATGAAGGTTTGCTGTGTTCTCCTTG-3’ and 5’-CTCGCCCTTGCTCACCTTTTTGCCTCCAACTGGTGGG-3’. The two PCR fragments were fused with an In-Fusion Cloning Kit to create pEpi1>Epi1::GFP. The standard methods of our construction are downloadable at the website (http://marinebio.nbrp.jp/ciona/). The official names of the constructs according to the nomenclature rule (Stolfi et al., 2015) are described in Table 1.

**Table 1.** Official names of vectors and transgenic lines reported in this study.

| Abbreviated name or aim of experiment in the manuscript | Full name according to the nomenclature rule |
| --- | --- |
| pSPCiEpi1>GFP | p*Ciinte*.REG.KY21.C1.13658444-13660489\|*Epi-1*>*GFP* |
| pEpi1>Epi1::GFP | p*Ciinte*.REG.KY21.C1.13658444-13660489\|*Epi-1*>*Ciinte.Epi-1::GFP* |
| pBSEF1α>TALEN::2A::mCherry | p*Ciinte*.REG.KY21.C14.1351654-1353597\|*Eef1a.a*>*TALEN::2A::mCherry* |
| pBSNut>TALEN::2A::mCherry | p*Ciinte*.REG.KY21.C14.1991592-1992604\|*Nut*>*TALEN::2A::mCherry* |
| *Epi-1* mutant lines | *Epi-1*<sup>TAL1</sup> and *Epi-1*<sup>TAL2</sup> |

### Electroporation

Unfertilized eggs were dechorionated with sodium thioglycolate and actinase E. Electroporation was done based on the previous reports (Corbo et al., 1997; Zeller, 2018). *Epi-1* knockout lines were created by the electroporation-mediated germ cell regeneration method (Yoshida et al., 2017) using pNut>Epi-1 TAL1::2A::mCherry and pNut>Epi-1 TAL2::2A::mCherry construct pairs.

### TALEN activity assays

The activities of TALENs were examined according to the previous reports (Treen et al., 2014; Yoshida and Treen, 2018). The *EF1α*>*TALEN::2A::mCherry* constructs were electroporated into 1-cell embryos. About 50 larvae that expressed mCherry in the entire body were picked up, and genomic DNA was isolated in bulk using a Wizard Genome DNA Isolation Kit (Promega). The target site of the corresponding TALENs was PCR amplified using ExTAQ DNA polymerase, Hot-Start version (Takara), and electrophoresed in 15% polyacrylamide gel according to the previous report (Ota et al., 2013). The primer sequences are 5’-ATCGATAAGCTTGATGCTTACTTTGAGAGTTTGAC-3’ and 5’-CTGCAGGAATTCGATCAGTCATACGTGTATATTAG-3’. The PCR fragments that exhibited shifted bands by the electrophoresis were subcloned into a cloning vector, and cloned fragments were sequenced to determine mutations. Detailed methods can be downloaded from the website (http://marinebio.nbrp.jp/ciona/).

### Genotyping

Genotyping was carried out according to the previous method (Sasakura et al., 2003). When a heterozygous mutant animal reached the reproductive stage, they were kept under dark conditions for 30 min, followed by light irradiation to promote spawning. *Ciona* is hermaphroditic, and the individuals are self-fertilize when eggs and sperm spawned from a single individual are mixed for a few hours. Fertilized eggs were collected in a 9-cm plastic dish and were incubated at 18℃ until the larval stage. When a portion of the larvae exhibited the sticky phenotype, a single larva was subjected to digestion in 50 μl of TE containing 0.2 μg/μl proteinase K for 3 h at 50℃, followed by 15 min at 95℃ to inactivate Proteinase K. 1 μl of the digested solution was used as the template of PCR. The PCR condition and primers were the same as those in the TALEN activity assay. The PCR fragments were subjected to 15% polyacrylamide gel electrophoresis and were cloned for sequencing.

### Scanning electron microscopy

Larvae were fixed in 0.1 M cacodylate and 0.1 M NaCl buffer (pH7.5) containing 1% glutaraldehyde and 3.7% formaldehyde for overnight at 4℃. The samples were post-fixed in 0.1 M cacodylate and 0.1 M NaCl buffer (pH7.5) containing 1% osmium tetroxide for 1 h at room temperature, followed by rinsing three times with 0.1 M cacodylate and 0.1 M NaCl buffer (pH7.5). The samples were dehydrated with a series of ethanol solutions (30, 80, 90, 95, 99.5, and 100%). In the 30 to 95% conditions, the dehydration was carried out on ice and incubated for 5 min. In the 99.5 and 100% conditions the dehydration was carried out at room temperature and incubated for 15 min. The samples were then rinsed three times with 100% t-butanol. The samples were placed onto filtered paper in an aluminum dish and then were incubated at −18℃ for 1 h. The frozen samples were dried with a freeze-drying device JFD-320 (JEOL) and GLD-420 (ULVAC). The dried samples were platinum-coated using JEC-3000FC (JEOL) and were subjected to observation under Neo Scope JCM-5000 (JEOL).

### Cellulose staining

Cellulose staining was carried out according to the previous report (Sasakura et al., 2005). Larvae were fixed in seawater containing 3.7% formaldehyde overnight at 4℃. After rinsing four times with PBST, the samples were incubated in PBST containing 1% goat serum for 1 h at room temperature to block non-specific staining. GFP-CBM3 (CZ00571, Nzytech) was suspended in 20 mM Tris-Cl pH 7.5, 20 mM NaCl, 5 mM CaCl2 according to the manufacturer’s instruction. The samples were overnight incubated at 4℃ in GFP-CBM solution 1/50 diluted in PBST containing 1% goat serum. After rinsing four times with PBST, the samples were incubated in anti-GFP antibody (04404-84, Nacalai Tesque) 1/200 diluted in PBST containing 1% goat serum for 1 h at room temperature. After rinsing four times with PBST and once with PBST containing 1% goat serum, the samples were incubated in Alexa 488 goat anti-rat IgG (Invitrogen) 1/200 diluted in PBST containing 1% goat serum overnight at 4℃. After rinsing four times with PBST, the samples were subjected to fluorescent microscopy.

### Epi-1 antibody, western blotting and immunohistochemistry

The polyclonal antiserum against Epi-1 that recognizes the fragment near the C-terminal region (N-RSGSTSPFGPEFPPVGGKK-C) was generated in rabbits by Cosmo Bio Co., Ltd. Approximately 200 larvae were dissolved in a 20 μl SDS sample buffer (2.6% SDS, 3.9% 2-Mercaptoethanol, 5.2% glycerol, 0.003% bromophenol blue and 81.4 mM Tris HCl, pH approx. 6.8). The samples were heated at 95℃ for 2 min, cooled on ice, and subjected to SDS-PAGE in a 10% polyacrylamide gel. For lysing larvae in the chorion, we added 0.02g glass beads (0.7-1.0 mm in diameter) in the SDS sample buffer. The samples were heated for 1 min at 95℃ and then vortexed for 1 min three times before electrophoresis. The separated proteins were transferred to an Immobilon membrane (Millipore). After staining with Coomassie brilliant blue, the membrane was blocked with PBST containing 7.5% non-fat milk for 1 h. The membrane was then incubated in anti-Epi-1 antibody 1/10,000 diluted in PBST containing 7.5% non-fat milk for 1 h. After washing with PBST four times at 10 min intervals, the membrane was incubated in HRP conjugated goat anti-rabbit IgG (H+L) antibody (Proteintech Group, Inc.) 1/20,000 diluted in PBST containing 7.5% non-fat milk for 1 h. After washing with PBST four times at 10 min intervals, the signal was detected with ECL plus (GE Healthcare) and X-ray film or a C-DiGit chemiluminescence scanner (Scrum).

### Alcian blue staining

Alcian blue staining was done according to the previous report (Sato and Morisawa, 1999). Larvae were fixed in the seawater containing 3.7% formaldehyde for 10 min at room temperature. After rinsing four times with PBST, the samples were incubated in Alcian blue staining solution pH 1.0 (Muto Pure Chemicals) for 15 min. After rinsing four times with PBST, the samples were subjected to microscopy.

### Staining of adhesives

The adhesive material was detected according to the previous report (Zeng et al., 2019). Larvae were fixed in the seawater containing 3.7% formaldehyde for 1 h at room temperature. After rinsing four times with PBST, the samples were incubated in PBST containing 5% fetal bovine serum for 1 h at room temperature. Then the samples were incubated overnight at 4℃ in 25 μg/ml biotinylated peanut agglutinin (PNA; Vector Laboratories) in PBST containing 5% fetal bovine serum. After rinsing six times with PBST, the samples were incubated in 1/300 Dylight488-conjugated streptavidin (Vector Laboratories) in PBST containing 5% fetal bovine serum for 1 h. After washing several times in PBST, the samples were subjected to microscopy.

---

**Supplementary Movie S1.** Swimming of a wild-type larva.

**Supplementary Movie S2.** Swimimng of a homozygous mutant larva in *Epi-1*. This larva adhered to the bottom of the plastic dish..

